# Bromelain Inhibits SARS-CoV-2 Infection in VeroE6 Cells

**DOI:** 10.1101/2020.09.16.297366

**Authors:** Satish Sagar, Ashok Kumar Rathinavel, William E. Lutz, Lucas R. Struble, Surender Khurana, Andy T Schnaubelt, Nitish Kumar Mishra, Chittibabu Guda, Mara J. Broadhurst, St. Patrick M. Reid, Kenneth W. Bayles, Gloria E. O. Borgstahl, Prakash Radhakrishnan

**Author notes:** **Correspondence:** Prakash Radhakrishnan, Eppley Institute for Research in Cancer and Allied Diseases, Fred & Pamela Buffett Cancer Center, University of Nebraska Medical Center, Omaha, NE, 68198-6805, USA.

## Abstract

Coronavirus disease 2019 (COVID-19) is caused by severe acute respiratory syndrome coronavirus-2 (SARS-CoV-2). The initial interaction between Transmembrane Serine Protease 2 (TMPRSS2) primed SARS-CoV-2 spike (S) protein and host cell receptor angiotensin-converting enzyme 2 (ACE-2) is a pre-requisite step for this novel coronavirus pathogenesis. Here, we expressed a GFP-tagged SARS-CoV-2 S-Ectodomain in Tni insect cells. That contained sialic acid-enriched N- and O-glycans. Surface resonance plasmon (SPR) and Luminex assay showed that the purified S-Ectodomain binding to human ACE-2 and immunoreactivity with COVID-19 positive samples. We demonstrate that bromelain (isolated from pineapple stem and used as a dietary supplement) treatment diminishes the expression of ACE-2 and TMPRSS2 in VeroE6 cells and dramatically lowers the expression of S-Ectodomain. Importantly, bromelain treatment reduced the interaction between S-Ectodomain and VeroE6 cells. Most importantly, bromelain treatment significantly diminished the SARS-CoV-2 infection in VeroE6 cells. Altogether, our results suggest that bromelain or bromelain rich pineapple stem may be used as an antiviral against COVID-19.

**Highlights:** 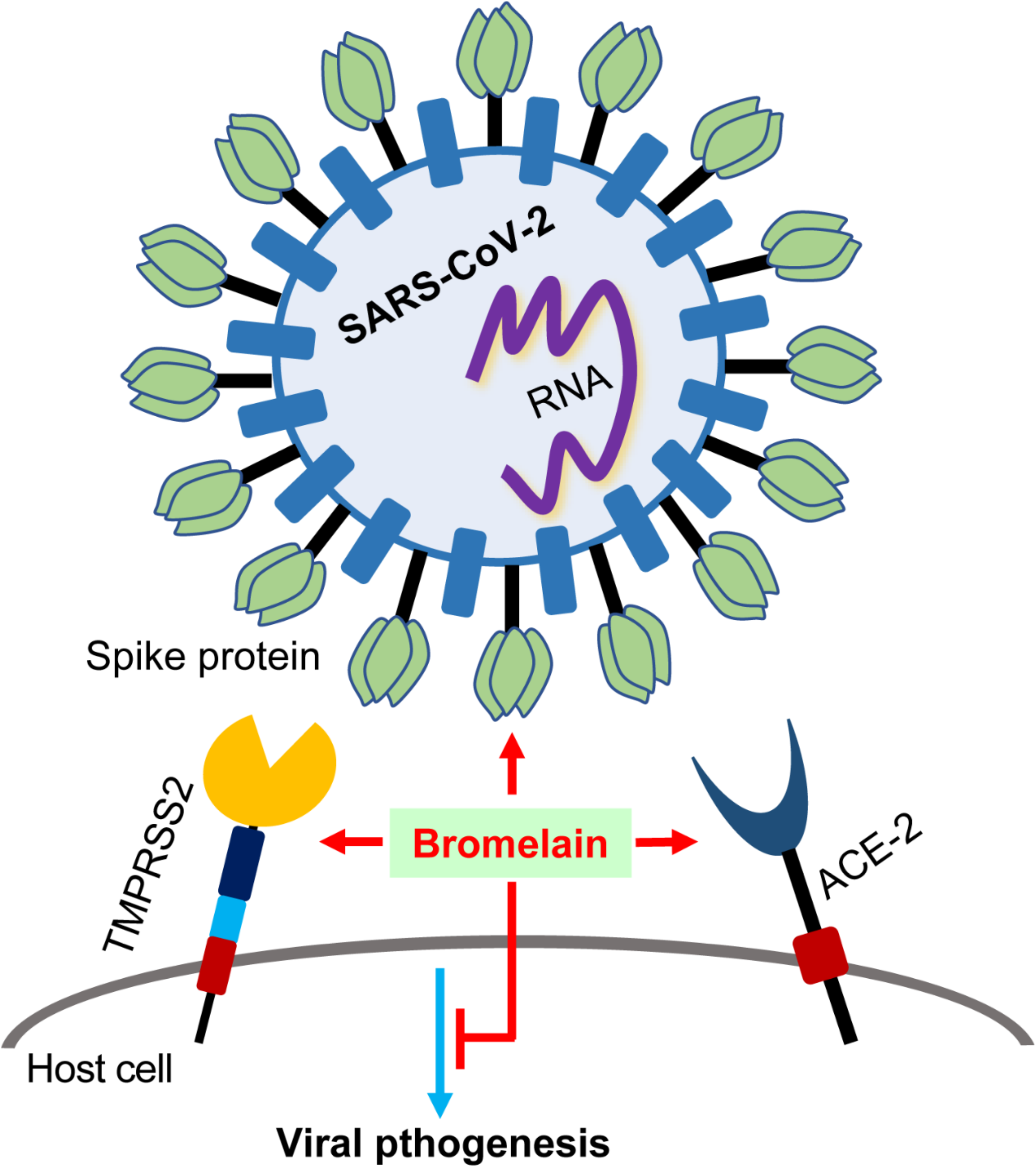

- Bromelain inhibits / cleaves the expression of ACE-2 and TMPRSS2
- Bromelain cleaves / degrades SARS-CoV-2 spike protein
- Bromelain inhibits S-Ectodomain binding and SARS-CoV-2 infection

## Introduction

The new coronavirus, severe acute respiratory syndrome coronavirus-2 (SARS-CoV-2) transmit rapidly from human-to-human resulted in ongoing pandemic. Older adults and people who have pre-existing conditions like heart disease, lung disease, diabetes, and cancers are at increased risk for developing severe complications from COVID-19 illness, including death. As of September 14, 2020, over 28,918,900 people have been infected worldwide with 922,252 deaths, of which 6,426,958 people in the US have been infected with 192,612 deaths (1). SARS-CoV-2 infects ACE-2 expressing lung, kidney, and testicular tissues of patients, which may lead to organ failure and sometimes death (2, 3). The SARS-CoV-2 homotrimeric viral spike protein (S1) binds to the host cell’s receptor ACE-2 for initial entry, followed by S2-mediated membrane fusion (4, 5). The S-protein undergoes proteolytic priming by TMPRSS2, which is expressed on the host cell surface (6). TMPRSS2 is a 70 kDa type II transmembrane glycoprotein with serine protease activity and expressed in many tissues (7). TMPRSS2 is predicted to have nine stabilizing disulfide bonds and two N-glycan sites (UniProtKB - O15393). TMPRSS2 priming activity is essential for many viral infections, including influenza A virus (8, 9), SARS-CoV (10), and SARS-CoV-2 (6). The SARS-CoV-2 Spike (S) protein is a heavily glycosylated transmembrane glycoprotein contains 1273 amino acids. It has several functional domains, including N-terminal domain (NTD), receptor-binding domain (RBD), fusion peptide (FP), heptad repeat 1(HR1), central helix (CH), connector domain (CD), transmembrane domain (TM), and a cytoplasmic tail (CT) (11). Recently, studies have demonstrated that SARS-CoV-2 spike protein has 22 N-linked and 2 O-linked glycan sites. Mostly these N-glycans are composed of oligomannose, hybrid and complex type, and the O-glycans are core 1 derived structures, which may play a significant role in protein folding, receptor binding, and immune escape (12–14).

Currently, COVID-19 patients are treated with a different antiviral (Favilavir and Remdesivir) (15–17), antimalarial (Chloroquine and Hydroxychloroquine) (18), viral protease inhibitors (Lopinavir and Darunavir) (18, 19), and anti-inflammatory agent (Tocilizumab) (20). However, the response rate is modest, and the safety and efficacy of those drugs against COVID-19 still needs to be further confirmed by clinical trials. Hence, there is an emergent need to repurpose existing drugs or develop new virus-based and host-based antivirals against SARS-CoV-2. Here, we explored the effect of bromelain on host cell receptor ACE-2, TMPRSS2, and SARS-CoV-2 S-Ectodomain. Bromelain is isolated from pineapple stem, a cysteine protease, and is used as a dietary supplement for treating patients with pain and inflammation (21) and thrombosis (22). Here, we show that bromelain inhibits SARS-CoV-2 infection in VeroE6 cells via altering the expression of ACE-2, TMPRSS2 and SARS-CoV-2 S-protein.

## Results and Discussion

### Bromelain inhibits the expression of ACE-2 and TMPRSS2

Recently, studies have shown that SARS-CoV-2 S-protein utilizes human ACE-2 as receptor for viral entry (4, 5). Hence, we explored the expression of ACE-2 and TMPRSS2 in various normal and cancerous cells. Of several normal and cancerous cells tested, VeroE6 and Calu-3 cells showed ACE-2 protein expression (Figure 1A), as well as basal level of TMPRSS2 protein (Figure 1B). It has been demonstrated that human ACE-2 is a type I transmembrane glycoprotein composed of 805 amino acids. It has an extracellular N-terminal domain that contains a zinc metallopeptidase domain (HEXH motif) and C-terminal domain (23, 24). ACE-2 has 7 potential N-glycosylation and 3 O-glycosylation sites, which may play a major role in protein folding and viral binding (13). The zinc metallopeptidase domain is responsible for binding to viral spike protein. ACE-2 extracellular domain has six cysteine residues with three stabilizing disulfide bonds (25). Additionally, TMPRSS2 is predicted to have 18 cysteine residues with nine stabilizing disulfide bonds in its extracellular domain (UniProtKB - O15393). Since ACE-2 and TMPRSS2 are enriched with cysteine residues that form disulfide bonds to stabilize the protein structure, we investigated the effect of bromelain (cysteine protease) on ACE-2 and TMPRSS2 expression. Bromelain reduces the expression of ACE-2 and TMPRSS2 in a dose-dependent (19, 37 and 75 μg/ml / 48 h) manner in VeroE6 cells (Figure 1C). Since bromelain took 48 hours to reduce the expression of ACE-2 in complete media, we investigated whether bromelain’s inhibitory effect is a direct or indirect action on these two proteins. For this, we treated VeroE6 cells with bromelain (75 ug/ml) in a shorter period (0-4 h) with serum-free conditions. Interestingly, bromelain reduced the expression of ACE-2 and TMPRSS2 at an early time-points onward (2 h) (Figure 1D). However, bromelain’s cysteine proteolytic activity was higher in ACE-2 than TMPRSS2. Next, we confirmed the bromelain’s activity by treating recombinant TMPRSS2 (Abnova, CA, USA) that showed a complete loss of TMPRSS2 (Figure 1E). We further validated these results by incubating VeroE6 cells with a cysteine protease inhibitor E-64 (4 μM) along with bromelain (75 μg/ml) and showed an intact ACE-2 and TMPRSS2 protein expression as compared to bromelain alone (Figure 1F). These results suggest that bromelain’s cysteine protease activity reduces the expression of ACE-2 and TMPRSS2 in VeroE6 cells. Since VeroE6 cells expressed ACE-2 and basal levels of TMPRSS2, and SARS-CoV-2 can infect VeroE6 cells (26, 27), we utilized this cell line for the rest of the studies.

**Figure 1.**
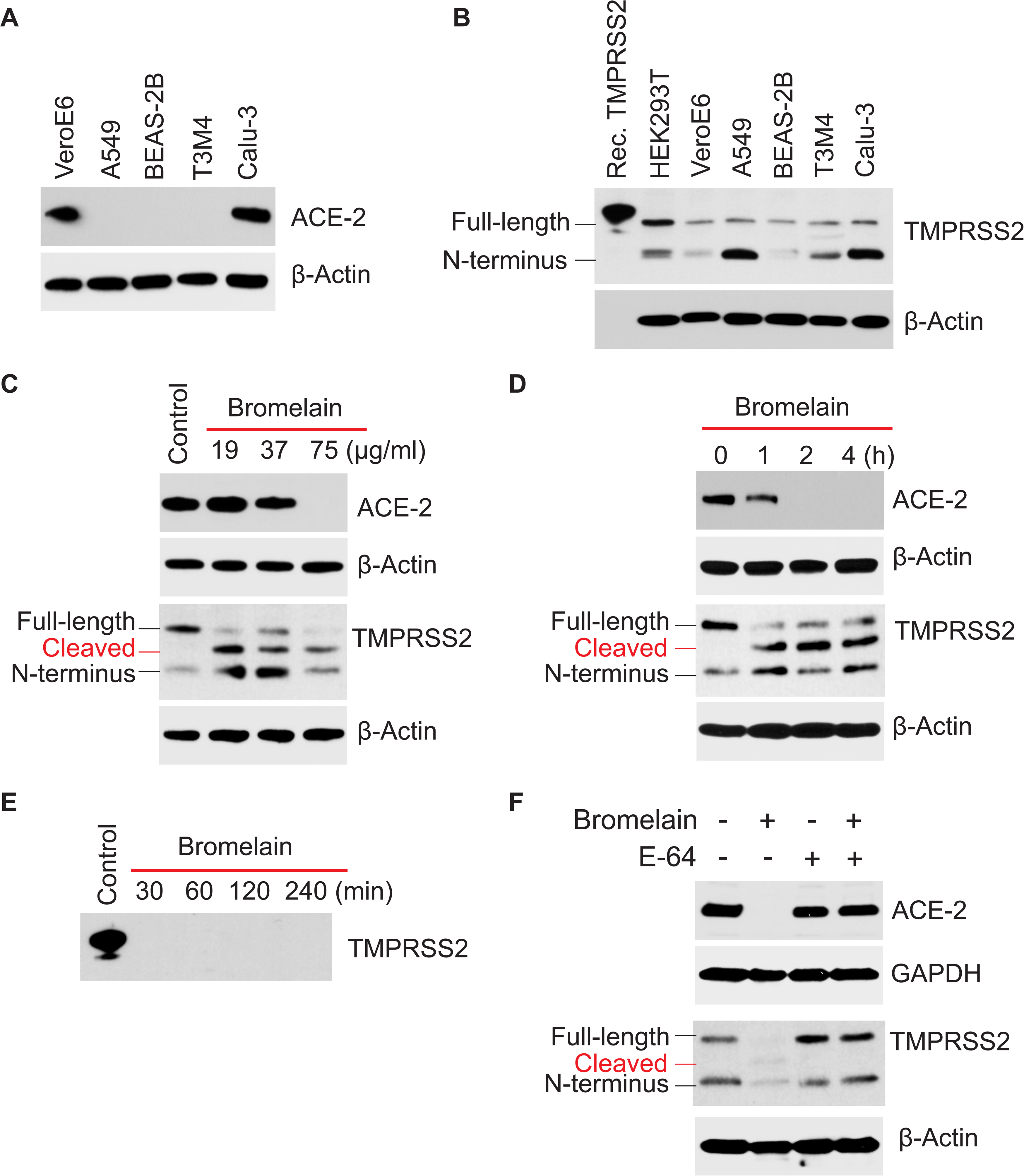
Bromelain inhibits ACE-2 and TMPRSS2 expression. **A** and **B** Immunoprobing of ACE-2 and TMPRSS2 in various normal and cancerous cells. **C.** Immunodetection of ACE-2 and TMPRSS2 in bromelain (19, 37, 75 μg/ml for 48 h) treated VeroE6 cells. **D.** Immunodetection of ACE-2 and TMPRSS2 in bromelain (75 μg/ml for 1-4 h) treated VeroE6 cells. **E**. Recombinant TMPRSS2 was treated with bromelain (1:1 ratio) for 30, 60, 120 and 240 min at 37°C. **F**. Immunoprobing of ACE-2 and TMRSS2 in bromelain (75 μg/ml) plus E-64 (4 μM) treated VeroE6 cells. Detection of β-actin served as a loading control.

### Bromelain cleaves SARS-CoV-2 Spike protein

Studies have shown that the SARS-CoV-2 spike protein has a 10 to 20-fold higher binding affinity with ACE-2 as compared to SARS-CoV-1 spike protein for viral pathogenesis (28). Hence, we cloned and expressed SARS-CoV-2 spike protein ectodomain that contains insect cell secretion signal (SP), NTD, RBD, FP, HR1, CH, and CD domains ((1-1220 amino acids) along with C-terminal GFP and 12X His tag and a mutated furin cleavage site (Figure 2A and Figure S1). The calculated molecular weight of the purified S-Ectodomain-GFP protein is ~165 kDa; however, we observed a higher molecular weight of S-Ectodomain (~215 kDa), which may be due to heavy N- and O-linked glycosylation (Figure 2B intent). Next, we determined the interaction between the purified S-Ectodomain and human recombinant ACE-2 by using surface plasmon resonance (SPR) technology. The SPR analysis revealed that S-Ectodomain binds with the receptor ACE-2. Further, the binding increased in a concentration-dependent manner and has a comparable binding affinity as control RBD binding to ACE-2 (Figure 2B). We also validated whether the expressed S-Ectodomain can be recognized by antibodies from COVID-19 patient samples; for this, we performed a Luminex serological assay with de-identified COVID-19 positive (n=6) and COVID-19 negative (n=6) UNMC patient samples. Luminex assay results showed a significantly increased median fluorescent intensity (MFI) of S-Ectodomain with COVID-19 positive patients’ samples (P<0.0001) (Figure 2C). These results indicate that the purified S-Ectodomain is correctly folded and active. Since S-Ectodomain has 22 N-glycosylation and 2 O-glycosylation sites (12–15), we explored the type of glycosylation on the purified S-Ectodomain. Figure 2D shows the schematic representation of the N- and O-linked glycosylation sites in S-protein (12, 13). Treatment of S-Ectodomain with N-glycanase (PNGase F) showed increased mobility shift and destabilization of the protein, which may be due to the loss of heavy N-glycans (Figure 2E). However, the treatment of S-Ectodomain with O-glycanase (specific for core 1 O-glycan) showed a slightly decreased protein mobility shift, which supports the previous findings that S-protein has only two O-linked core 1 derived glycans in its RBD backbone (Figure 2E). Conversely, treatment of S-Ectodomain with Sialidase A (which removes sialic acid group) either alone or in combination with either N-glycanase or O-glycanase or together induced a noticeable change in the migration of S-Ectodomain. This decreased mobility shift of S-Ectodomain may be due to the loss of negatively charged sialic acid groups in the N- and O-linked glycans (Figure 2E). These results indicate that the SARS-CoV-2 Spike protein has both highly sialylated N- and O-linked glycans.

**Figure 2.**
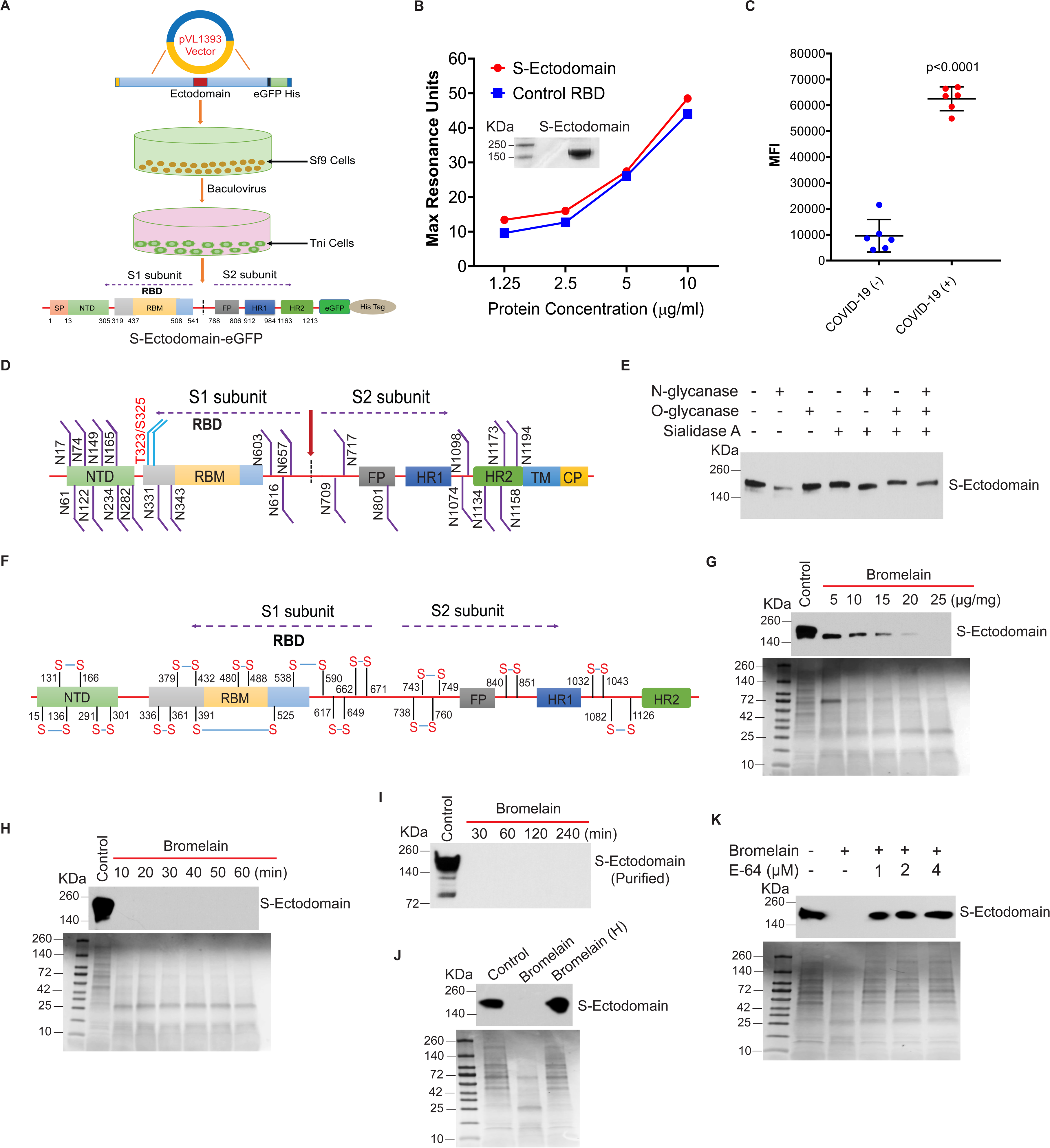
Bromelain cleaves SARS-CoV-2 S-Ectodomain. **A**. Schema of SARS-CoV-2 S-Ectodomain-GFP-His tag protein expression. **B**. SPR Max Resonance Units (RSU) at equilibrium as a function of the concentration of control RBD and purified S-Ectodomain using immobilized ACE-2. Purified S-Ectodomain-GFP-His in Tni insect cells showed ~215 kDa molecular weight (intent). **C**. Luminex assay (Median Fluorescent Intensity) of S-Ectodomain with COVID-19 positive (n=6) and negative (n=6) patients’ samples. **D**. Schematic representation of SARS-CoV-2 S protein’s N- and O-linked glycosylation sites. **E**. Immunoblotting analysis of Sialidase A, O-glycanase and N-glycanase treated S-Ectodomain. **F**. Schematic representation of SARS-CoV-2 S-protein with predicted cysteine amino acids position and disulfide bridges. **G** and **H** Immunoblotting of S-Ectodomain in bromelain treated Tni cell supernatant varying dose- (5, 10, 15, 20, and 25 μg/mg of total protein) and time-(25 μg/mg of total protein for 0 to 60 min). **I**. Immunoprobing of bromelain treated purified S-Ectodomain (1:10 ratio) with 30, 60, 120, and 240 min. **J.** Immunoprobing of S-Ectodomain treated with heat-inactivated bromelain (80°C/8 min). **K.** Detection of S-Ectodomain in bromelain (25 μg/mg of total protein) plus E-64 (1, 2, and 4μM) treated Tni cell supernatant. SimplyBlue stained gel images of bromelain treated Tni cell supernatant served as a loading control. Experiments were performed thrice, and a representative image is presented.

The receptor-binding domain has five antiparallel β sheets (β1, β2, β3, β4 and β7) with short connecting helices and loops, which form the core. Within this core structure, there is an extended insertion like motifs called receptor binding motif (RBM), which has contacting residues of coronavirus that bind to ACE-2 (29). SARS-CoV-2 S-protein has 30 cysteine amino acids with 15 stabilizing disulfide bonds in the Ectodomain (UniProtKB: P0DTC2) (Figure 2F). The RBD domain alone has nine Cysteine residues, eight of which form four disulfide linkages. Out of these four pairs, three are in the core (Cys336– Cys361, Cys379–Cys432 and Cys391–Cys525), which help to stabilize the β sheet structure. The remaining one pair (Cys480–Cys488) connects the loops in the distal end of the RBM (29). Since bromelain has cysteine protease activity, we examined the effect of bromelain on S-Ectodomain expression. Incubation of Tni insect cell supernatant containing S-Ectodomain with bromelain showed reduced expression of S-Ectodomain in a dose-dependent manner (5, 10, 15, 20 and 25 μg/mg of total protein/hour) (Figure 2G). Next, we investigated bromelain induced cleavage of S-Ectodomain in a time-dependent manner. Bromelain treatment (25 μg/mg of total protein) completely cleaved S-Ectodomain as early as 10 min at 37°C (Figure 2H). However, a lower concentration of bromelain (10 μg/mg of total protein) partially cleaved S-Ectodomain even up to 120 min at 37°C (data not shown). Next, we confirmed these results by treating the purified S-Ectodomain with bromelain (1 μg/10 μg purified protein) shows a complete cleavage of the S-Ectodomain in a time-dependent manner (Figure 2I). To further validate these results, we used heat-inactivated bromelain on Tni supernatant that showed an intact S-Ectodomain as compared to bromelain (25 μg/mg of total protein) treatment (Figure 2J), also we have incubated S-Ectodomain expressing Tni cells supernatant with E-64 (cysteine protease inhibitor, 1, 2 and 4 μM) plus bromelain (25 μg/mg of total protein) for 30 min, which showed an intact S-Ectodomain as compared to bromelain alone treatment (Figure 2K). Altogether, these results demonstrate that bromelain can act as a double-edged sword to degrade both host-derived ACE-2, TMPRSS2, and viral derived S-Ectodomain and thus may be predicted to inhibit SARS-CoV-2 infection.

### Bromelain inhibits S-Ectodomain binding and SARS-CoV-2 infection

Since bromelain digested ACE-2, TMPRSS2, and S-Ectodomain, we investigated the effect of bromelain on the interaction between S-Ectodomain and VeroE6 cells by immunofluorescence. Interestingly, we observed a reduced binding of S-Ectodomain in bromelain (75 μg/ml for 1 h) treated VeroE6 cells as compared to vehicle control (Figure 3A). We further confirmed these results by flow cytometry. We detected a decreased binding of S-Ectodomain in bromelain pre-treated VeroE6 cells as compared to the vehicle (Figure 3B, histogram). The mean fluorescent (GFP) intensity of S-Ectodomain binding was significantly reduced in bromelain treated VeroE6 cells as compared to vehicle (p<0.0001) (Figure 3C). To further validate these results, we incubated E-64 (cysteine protease inhibitor) plus bromelain and S-Ectodomain together with VeroE6 cells, which resulted in enhanced binding of S-Ectodomain to VeroE6 cells as compared to bromelain alone (Fig. 3D). These results demonstrate that bromelain can act as a double-edged sword by cleaving both ACE-2 and S-Ectodomain and thereby inhibit S-protein binding in VeroE6 cells. Since bromelain inhibits S-Ectodomain binding to VeroE6 cells, we investigated the possibility of bromelain can inhibit SARS-CoV-2 infection in VeroE6 cells. To achieve this, we have infected vehicle control and bromelain pre-treated VeroE6 cells with SARS-CoV-2 virus (BEI_USA-WA1/2020) titer of 0.01 MOI. The infected cells were fixed and stained with SARS-CoV-2 S-protein specific antibody, which showed a reduction in the number of infected cells in bromelain treatment (Figure 3E, representative image from n=6). The percentage of green fluorescent positive cells was quantified by using Nikon D-Elements software. We observed a significantly decreased percentage of SARS-CoV-2 infected VeroE6 cells in bromelain treatment as compared to the vehicle (p=0.0011) (Figure 3E). Taken together, our results indicate that that bromelain can effectively inhibit SARS-CoV-2 infection in VeroE6.

**Figure 3.**
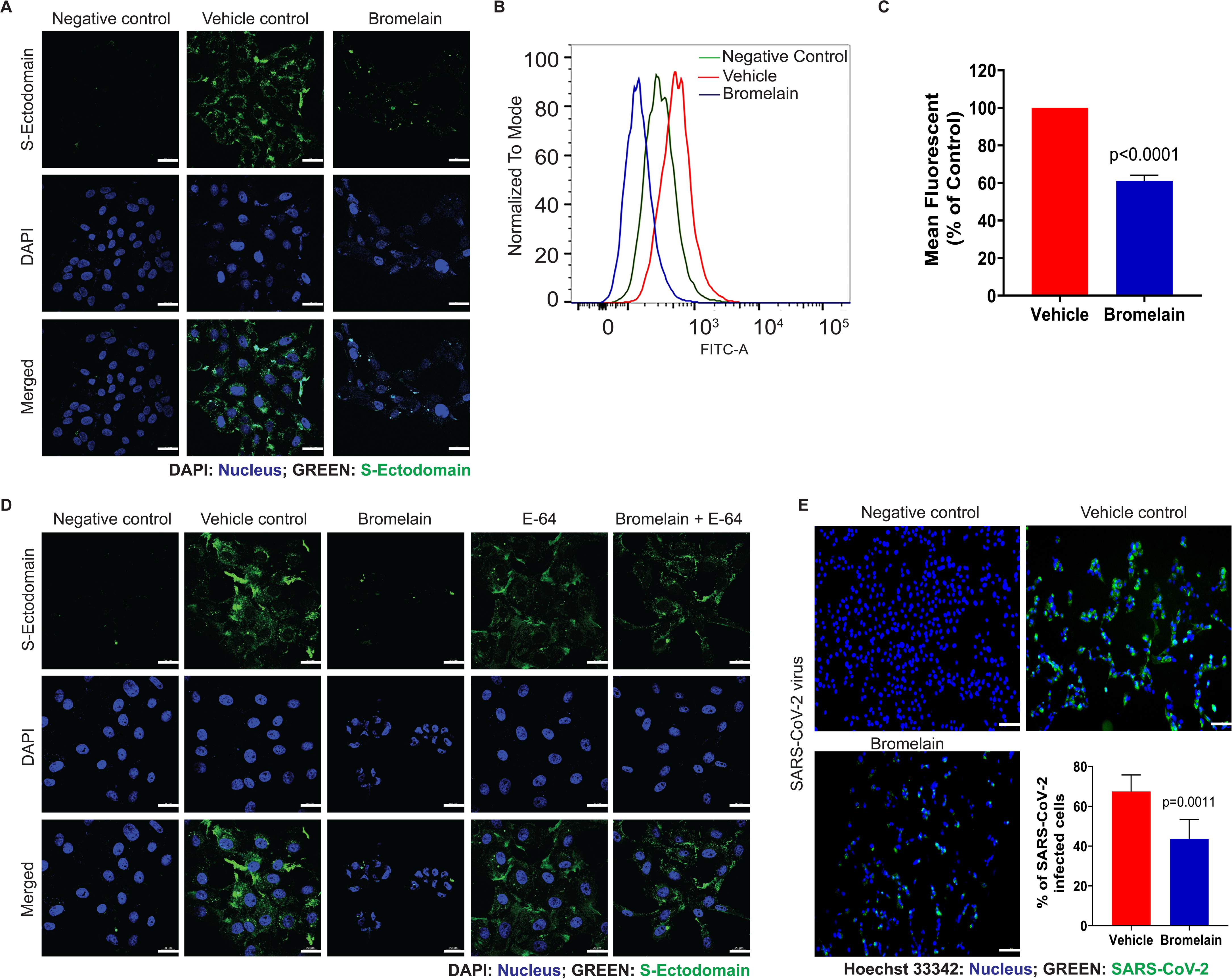
Bromelain inhibits SARS-CoV-2 infection in VeroE6 cells. **A**. Immunofluorescence analysis of SARS-CoV-2 S-Ectodomain binding in vehicle control and bromelain (75 μg/ml /2h) pre-treated VeroE6 cells (n=3). Incubation of VeroE6 cells with GFP-His tag protein served as a negative control. Nuclei were stained with DAPI. Scale bar = 20 μm. **B**. Flow cytometry analysis of S-Ectodomain binding in-vehicle control and bromelain (75μg/ml /2h) pre-treated VeroE6 cells (n=3). **C**. The mean fluorescent intensity was measured by using FlowJo software. **D**. Immunofluorescence of S-Ectodomain binding in vehicle, bromelain (75 μg/ml) plus E-64 (4 μM) treated VeroE6 cells (n=3) for 2 h and analyzed as described above. **E**. SARS-CoV-2 (BEI_USA-WA1/2020) infection in vehicle and bromelain treated (75 μg/ml / 2h) VeroE6 cells (n=6). The percentage of SARS-CoV-2 infected cells in each well was quantified by using Nikon NIS-Elements D software. Scale bar = 100 μm. Nuclei were stained with Hoechst 33342. Mean ± SD (n=6). P < 0.05 was considered statistically significant.

Recently, studies have demonstrated that SARS-CoV-2 S-protein has high homology among other coronaviruses (76% identity with SARS-CoV) (30) with conserved cysteine amino acids (UniProtKB: P59594). This indicates that bromelain may be used as a broad antiviral agent against SARS-CoV2 and other related family members.

Currently used drugs against SARS-CoV-2 has potential side effects. Vaccine trials have started against COVID-19, but a vaccine is not expected to be ready before the end of 2020 for human use. Further, the host immune response against SARS-CoV-2 is not fully understood. It differs between individuals, and also re-infection of individuals with SARS-CoV-2. Clinical trials are ongoing with COVID-19 patients by individually targeting either ACE-2, TMPRSS2, or Spike protein. For the first time, our results demonstrate that bromelain can inhibit SARS-CoV-2 infection by targeting all three host ACE-2 and TMPRSS2, and SARS-CoV-2 S-proteins. Also, studies have shown that bromelain can be used to treat patients with pain and inflammation, and is well absorbed and remains biologically active (21, 31). Therefore, either bromelain or bromelain rich pineapple stem could be used as an antiviral for treating COVID-19 and future outbreaks of other coronaviruses.

## Materials and Methods

### Cells and cell culture conditions

The African green monkey kidney epithelial cells (VeroE6, ATCC, VA, USA), human lung adenocarcinoma cells (A549, a kind gift from Maher Abdulla, Department of Pathology and Microbiology, UNMC and Calu-3, a kind gift from John Dickinson, Internal Medicine Division of Pulmonary, Critical Care and Sleep, UNMC), human normal bronchial epithelial cells (BEAS-2B, a kind gift from Todd Wyatt, Internal Medicine Division of Pulmonary, Critical Care and Sleep, UNMC), human Pancreatic cancer cells (T3M4, a kind gift from Michael A Hollingsworth, Eppley Institute for Research in Cancer, UNMC), and human embryonic kidney epithelial cells (HEK293T, ATCC, VA, USA) were used for this study. VeroE6, A549, T3M4, and HEK293T were grown in DMEM (Gibco, MA, USA) with 10% Fetal bovine serum (FBS, Corning, MA, USA) and 1X penicillin and streptomycin (Corning, MA, USA). BEAS-2B cells were grown in DMEM with 5% FBS and 1X penicillin and streptomycin. All the cells were maintained at 37°C with 5% CO2 in a humidified chamber.

### Recombinant S-Ectodomain-GFP baculovirus preparation

The full-length genome sequences of 45 SARS-CoV-2 isolates were analyzed. Bat Coronavirus genome annotations were used to map the spike protein on the Wuhan CoVs. Multiple sequence alignment for 45 spike proteins was carried out, and the full-length consensus sequence was derived using the Gene Calc algorithm. The membrane-spanning region was identified using DNAstar Protean software. The cDNA for SARS-CoV-2 Spike protein ectodomain residues 1 to 1220 (S-Ectodomain-GFP) was chemically synthesized with optimal insect cell codons (Genscript USA Inc, NJ, USA) and cloned into pVL1393 (Expression Systems, CA, USA) with an insect cell secretion signal. This construct includes complete S1 and S2 regions and ends just before the transmembrane region (Figure S1). The furin cleavage site is mutated from RRAR to GSAS (28). Tobacco Etch Virus (TEV) protease cleavage site was added to the C-terminus, followed by a flexible linker, eGFP, and 12X histidine tag. A recombinant baculovirus (P0) was produced in SF9 cells (Expression Systems, CA, USA) as described in the protocol of the BestBac^tm^ 2.0 Δ v-cath/ChiA Baculovirus Cotransfection Kit (Expression Systems, CA, USA). P1 baculovirus was generated with suspended Sf9 cells (2 ×10^6^ cells/ml) with a multiplicity of infection (MOI: number of virus particles/number of cells) of 0.1. Viral titer was measured using a flow cytometric assay for gp64 expression (Expression Systems, CA, USA).

### Recombinant S-Ectodomain-GFP protein expression and purification

Tni cells (2×10^6^ cells/ml) suspended in 1L of ESF-AF media at were infected with the P1 virus at an MOI of 5. Cells were harvested 72 hours post-infection by centrifugation (500xg, 15 minutes). One cOmplete™ULTRA protease inhibitor tablet (Roche, MA, USA) per liter was added before centrifugation. The supernatant was concentrated by tangential flow filtration (TFF) using a Pellicon^®^ 2 mini-cassette system (Millipore, MO, USA) with a 30 kDa pore size. The concentrated supernatant was adjusted to 40 mM TRIS-HCl and 1 M sodium chloride. Then the pH was adjusted to 8.0 with sodium hydroxide and was then centrifuged (14,500xg, 30 minutes) and sterile filtered using a Thermo Scientific™Nalgene™Rapid-Flow™90 mM Filter Unit-1000 ml. For affinity purification, 10 ml of Ni Sepharose excel resin (Cytiva life sciences, USA) was added for every 1 L of culture (pre-TFF volume) and incubated overnight, gently stirring at 4°C. Stirring was stopped to allow the resin to settle for 1 hour, and then the supernatant was decanted, the remaining resin and media were mixed well and poured into a gravity flow column (Bio-rad, Hercules, CA, USA). The column was treated with 3 column volumes (CVs) of wash buffer (40 mM TRIS-HCl pH 8 with 1M sodium chloride), 3 CV of wash buffer with 20 mM imidazole added, 3 CV of wash buffer with 40 mM imidazole, and finally 3 CV of wash buffer with 60 mM imidazole before protein elution with 3 CV of wash buffer with 500 mM imidazole. S-Ectodomain-GFP protein was concentrated with an Amicon^®^ Ultra 15 Centrifugal Filter (100 kDa MWCO). It was injected onto a HiLoad™16/600 Superdex™200 prep grade FPLC size-exclusion column using 1x PBS (Fisher scientific, MA, USA) as a running buffer. The peaks were collected with 1 mL fractions. Pure fractions were identified by SDS-PAGE (MW ~250 kDa), pooled, and concentrated to 0.5 mg/ml.

### Surface Plasmon Resonance (SPR) based hACE2 binding assay

The recombinant hACE2-AviTag protein (Acro Biosystems, DE, USA) was captured on a sensor chip in the test flow channels. Samples of 300 μl of freshly prepared serial dilutions of the purified recombinant protein were injected at a flow rate of 50 μl/min (contact duration 180 seconds) for the association. Responses from the protein surface were corrected for the response from a mock surface and for responses from a buffer-only injection. Total hACE2 binding and data analysis were calculated with Bio-Rad ProteOn Manager software (version 3.1).

### Luminex serological assay

This assay was performed using the purified SARS-CoV-2 S-Ectodomain (5 μg) coupled to the surface of group A, region 43, MagPlex^®^ Microspheres (Luminex Corp, IL, USA). Microsphere coupling was performed using the Luminex xMAP Antibody Coupling Kit (Luminex Corp, IL, USA) according to the manufacturer’s instructions. The protein-coupled microspheres were re-suspended in PBS-TBN buffer (1X PBS containing 0.1% Tween 20, 0.5% BSA and 0.1% sodium azide) at a final stock concentration of 2×10^6^ microspheres per ml, with 2.5×10^3^ microspheres used per reaction. The serum samples were obtained from de-identified COVID-19 patients (n=6) and healthy donors (n=6) at UNMC. Fifty microliters of each serum sample (diluted 1:25 with 1X PBS-TBN buffer) was mixed with 50 μl of the S-Ectodomain coupled microspheres in a 96 well-plate (Greiner Bio-One). The assay plate was incubated for 30 minutes at 37°C with shaking at 700 RPM and then washed 5 times with 1X PBS-TBN buffer. Then, the plate was incubated with biotin conjugated goat anti-human IgG (Abcam, MA, USA), labeled with streptavidin R-phycoerythrin reporter (Luminex xTAG^®^ SA-PE G75) for 1 h at 25°C with shaking at 700RPM. Then, the plate was washed 5 times with 1X PBS-TBN buffer. Finally, the plate was re-suspended in 100 μl of 1X PBS-TBN buffer and incubated for 10 minutes at 25°C with shaking at 700 RPM. The microplate was assayed on a Luminex MAGPIX™System, and results were reported as median fluorescent intensity (MFI).

### Bromelain treatment

VeroE6 cells were grown in 10% FBS containing media and treated with bromelain (Sigma-Aldrich, MA, USA) at different concentrations (19, 37, and 75 μg/ml) for 48 h. For the time-dependent study, the aforementioned cells were treated with bromelain (75 μg/ml) in a serum-free condition for 0-4 h. After the end of the treatment, the cells were washed and lysed with RIPA lysis buffer (ThermoFisher, MA, USA) containing protease inhibitors (Roche, MA, USA). SARS-CoV-2 S-Ectodomain expressing Tni cell media were treated with different concentrations of bromelain (5, 10, 15, 20, and 25 μg) for 1 mg of total protein at 37°C for 1h. Recombinant TMPRSS2 (Abnova, CA, USA) was treated with bromelain (1:1 ratio) at 37°C for different time points (30, 60, 120, and 240 mins). For the time-dependent study, Tni media were treated with 25 μg of bromelain/mg of total protein at 37°C for different time points (30, 60, 120, and 240 mins). For bromelain heat inactivation assay, bromelain was heated at 80°C for 8 min and then treated with S-Ectodomain expressing Tni cell supernatant (25 μg of bromelain/mg of total protein) at 37°C for 30 min. For cysteine protease inhibitor (E64) treatment assay, bromelain plus E-64 (1, 2, and 4 μM) was used to incubate with S-Ectodomain expressing Tni cells supernatant (25 μg of bromelain/mg of total protein) at 37°C for 30 min. For treatment with VeroE6 cells, bromelain (75 μg/ml) plus E-64 (4μM) was incubated with VeroE6 cells at 37°C for 2 h. At the end of the reaction, 4X laemmli buffer was added in the mixture and boiled at 100°C for 5 min. The protein concentration in the cell lysate and media was measured by BCA kit (ThermoFisher, MA, USA)as per the manufacturer’s instructions.

### SDS-PAGE and Western blot analysis

Equal concentrations of media were mixed with 4X loading buffer (Invitrogen, MA, USA) and boiled at 100°C for 5 min. The SDS-PAGE was performed in 4-20% gradient gel (Bio-Rad, CA, USA). We used SimplyBlue SafeStain (Invitrogen, MA, USA) to visualize the protein band in SDS-PAGE gel. The gel image was captured using the ChemiDoc Imaging System (Bio-Rad, CA, USA). For western blot, equal concentrations of cell lysate and media proteins were transfer into 0.45 μm PVDF membrane (Millipore, MA, USA). Further, blocked with 5% skimmed milk powder at room temperature for 1 h, the membrane was incubated with the target primary antibody with desired dilution (anti-ACE-2, 1:1000 (Cell signaling technology, MA, USA); TMPRSS2, 1:1000 (Santa Cruz Biotechnology, TX, USA); anti-SARS Spike protein 0.05 μg/ml, (Novus Biologicals, CO, USA) at 4°C for overnight. For loading, control membranes were probed with anti-β-actin and anti-GAPDH antibodies (1:5000, Cell signaling technology, MA, USA). The membrane was washed with 1X TBST (3 × 5 min) and then incubated with respective secondary antibodies (Horse anti-mice 1:2000; Horse anti-rabbit 1:2000, (Cell signaling technology, MA, USA) and developed by using ECL chemiluminescent reagent (Bio-Rad, CA, USA).

### Deglycosylation of the SARS-CoV-2 S-Ectodomain

Removal of N-glycans, O-glycans, and Sialic acids from the purified S-Ectodomain was performed by using a Deglycosylation kit (Agilent, CA, USA) as per the manufacturer instructions. Briefly, 1 μg of S-Ectodomain was used for N-glycanase, O-glycanase, and Sialidase A (1μl/reaction) treatment at 37°C for 3 h. After treatment, the protein-enzyme mixtures were mixed with 2x Laemmli buffer and boiled for 100°C for 5 min and then immunoprobed with anti-Spike protein-specific antibody as described above.

### Spike protein binding assay

#### Confocal microscopy

VeroE6 (1 × 10^5^/ well) cells were seeded on coverslips in a 12 well plate. After 24 h, the cells were treated with 75 μg/ml of bromelain and mock (1XPBS) for 1 h in serum-free media. Next, the cells were washed with 1X PBS two times and incubated with SARS-CoV-2 S-Ectodomain-GFP (1 μg) in a serum-free media for 2 h. Cells were washed and fixed with 4% paraformaldehyde for 15 min. The Spike protein binding was analyzed by using a Zeiss LSM 800 confocal laser scanning microscope (UNMC Core facility). Incubation of cells with GFP-His tag protein (Sino Biological, PA, USA) served as a negative control.

#### Flow cytometry

VeroE6 cells were treated with 75 μg/ml of bromelain and mock (1XPBS) for 1 h in serum-free media. Next, the cells were detached by using 2mM EDTA in PBS and washed twice with 1X PBS. 1 × 10^6^ cells were incubated with S-Ectodomain-GFP (1 μg) in a serum-free media for 2 h. The cells were washed and resuspended in HBSS media for flow cytometry analysis at the UNMC core facility (BD LSR II flow cytometer). The results were analyzed by FlowJo software.

### SARS-CoV-2 infection assay

VeroE6 (1×10^4^ cells) were seeded in 96 well plate. After 24 h, cells were washed and treated with bromelain in a serum-free medium (n=6). The incubation of cells with PBS served as a negative control (n=6). After 2 h of treatment, cells were washed and replaced with serum-free media for infection with SARS-CoV-2 (strain: BEI_USA-WA1/2020) multiplicity of infection (MOI) of 0.01 for 1 h at 37°C. After infection, cells were washed and replaced with 5% FBS containing media. After 24 h post-infection, cells were fixed with 4% buffered paraformaldehyde (Electron Microscopy Sciences) for 15 min at room temperature. The fixed cells were washed with PBS then permeabilized in 0.1% Triton X100 PBS solution for 15 min then blocked in 3% BSA PBS solution. The cells were incubated with anti-S protein Rab (Sino Biological, PA, USA) at 1:1000 in the blocking solution overnight at 4°C, followed by incubation with 1:2000 diluted Alexa Fluor 488 conjugated secondary antibody (Thermo Fisher, MA, USA) for 1 h at room temperature. The cells were counterstained for nuclei with Hoechst 33342 (Thermo Fisher, MA, USA). The fluorescent images were captured by using a Nikon Eclipse Ts2R fluorescent microscope. Total and virus-infected cells were counted by using Nikon NIS-Elements D software.

### Statistical Analysis

Statistical analysis was performed with GraphPad Prism version 8 using a two-tailed unpaired *t*-test. Statistical significance was determined at *P* <0.05.

## Acknowledgments

We thank Dr. Kenneth Cowan (Fred and Pamela Buffett Cancer Center, UNMC) for his strong support and Elizabeth Coyle (FDA, USA) for technical assistance in SPR analysis. We thank Dalia ElGamal (Eppley Institute for Research in Cancer, UNMC) for her help. We thank all the staff members of Nebraska Medicine who care for COVID patients. The authors on this manuscript are supported in part by the National Cancer Institute (NIH) grant (R01CA208108-01A1), the Fred and Pamela Buffett Cancer Center start-up funds (to P.R), and the Fred and Pamela Buffett NCI Cancer Center Support Grant (P30CA036727) (to G.E.O.B).

## Author contributions

P.R. conceived the idea; S.K, G.E.O.B., and P.R. designed research; S.S, A.K.R, W.E.L, L.R.S, S.P.M.R., and M.J.B. performed research; B.G., and K.W.B. contributed new reagents/analytic tools; S.S, A.K.R, G.E.O.B, S.P.M.R., and P.R. analyzed data; G.E.O.B., and P.R. wrote the paper.

## Supplementary Figures

**Figure S1.** The complete amino acid sequence of purified insect cell S-Ectodomain-GFP from the original Wuhan strain of SARS-CoV-2. Furin mutation is highlighted in yellow. The TEV site (cyan letters), flexible region (red), eGFP (green), and 12X-His tag (purple) are indicated.

